# Complete360: A Multi-Omics Liquid Biopsy Platform with Attomole Sensitivity and High Reproducibility

**DOI:** 10.1101/2025.05.16.654403

**Authors:** Raghothama Chaerkady, Liang Zhao, Anirudh Kashyap, Morgan Fair, Qing Wang

## Abstract

Advancements in liquid biopsy platforms can uncovered disease-associated molecular changes that may indicate disease. Despite recent progress in the detection of a number of analytes, the current liquid biopsy platforms struggle with specificity, sensitivity, and reproducibility, limiting their clinical utility and high-throughput applicability. Here, we describe an ultra-sensitive mass spectrometry-based liquid biopsy platform, *Complete360*, with an optimized library of detection parameters, meticulously refined assays to enable the simultaneous detection and quantification of a broad range of body-fluid proteins, metabolites, and lipids. The system addresses sample-specific and target-specific noises, as well as the micro-environments of proteomics during co-elution, achieving unprecedented levels of reproducibility, specificity, and sensitivity. This platform has facilitated the detection and quantification of numerous multi-omics signatures in human plasma samples, uncovering novel disease mechanisms never identified by previous approaches. Our study establishes a platform for rapid discovery and validation of molecular changes in disease panels, demonstrating its potential utilities in applications meeting clinical regulation guidelines.

## Introduction

Blood is one of the most accessible clinical samples for liquid biopsy, making it invaluable for molecular diagnostics in human disease. Several methods have been previously reported for detecting and quantifying disease associated molecular changes in blood, particularly nucleic acids in the context of human cancers, and some of these methods have successfully led to commercial applications in clinical settings^1–3^. Some disease development such as certain cancers is often associated with genetic alterations such as mutations and translocations, these changes can be measured from circulating DNA in the blood^4^. However, many human diseases—such as cardiovascular, neurodegenerative, some cancer, and autoimmune disorders—do not involve genomic alterations and cannot be assessed through sequencing methods. Additionally, genetic predispositions to cancers, once measured in a person to assess cancer risks, the subsequent measurement provides no additional information for clinical intervention, which limits the clinical utilities for sequencing technologies aimed at detecting nucleic acid changes in disease management.

In contrast, proteins and small molecules (metabolites and lipids) in human blood serve as excellent surrogates that reflect real-time changes in health and disease status^5, 6^. Virtually if all human diseases could be monitored through variations in protein or small molecule surrogate levels in blood, most clinical tests can be achieved by measuring proteins or small molecules^7^. However, accurately measuring these molecules in blood has proven to be a significant challenge, limiting their clinical applications.

Traditional methods for measuring proteins and metabolites in blood can be categorized into two main approaches. The first is the affinity-based method. The second approach involves mass spectrometry, which can be performed in either targeted or untargeted modes. The advent of advanced mass spectrometers, such as the timsTOF HT from Bruker and Astral from ThermoFisher, in combination with low abundant blood protein enrichment strategies, has allowed researchers to detect ∼6,000 proteins from plasma samples using untargeted mass spectrometry ^8, 9^. While untargeted mass spectrometry provides a broader range of identified proteins or small molecules, it generally lacks sensitivity and/or specificity. Using targeted proteomics, we achieved highly sensitive detection of specific mutated forms of proteins as early-stage cancer-specific protein changes, and extremely low abundant neoantigens through modified targeted approaches, demonstrating high sensitivity after a series of parameter optimizations aimed at ultra-low abundance detection^6, 10–13^.

We hypothesized that if we could recursively optimize the detection for each protein individually and compile these methods into a single assay, we could ensure accurate detection and quantification across all human blood proteins, achieving the sensitivity and reproducibility required for clinical applications. By optimizing liquid chromatography-mass spectrometry (LC-MS) parameters and developing customized sample preparation procedures for each of all specific protein targets, we reached unprecedented sensitivity for each protein and small molecule target. Here, we introduce *Complete360*, a multi-omics platform designed to advance both research and clinical investigations, capable of detecting over 10,000 human proteins and more than 2,000 metabolites and lipids in blood samples. This platform operates through a fully optimized and automated process that integrates sample preparation, mass spectrometry method clusters, and data analysis. *Complete360* addresses the limitations of traditional mass spectrometry by significantly enhancing the reproducibility and detectability of proteins, metabolites, and lipids often missed by other technologies. It provides more accurate multi-omics data by capturing higher magnitudes of biological differences between diseased bloods and controls. This practicality ensures its feasibility for real-world clinical applications.

## Results

### *Complete360* Platform: Integrating Extensive Data Acquisition and Curation using Artificial Intelligence (AI)

The *Complete360* platform comprises of 1) CompleteBank development for targeted assay panel construction, 2) extraction of plasma proteins, metabolites, and lipids from blood samples, 3) mass spectrometric analysis, and 4) data analysis using CompletePeaking Algorithm for report (Figure 1a).

**Figure 1.**
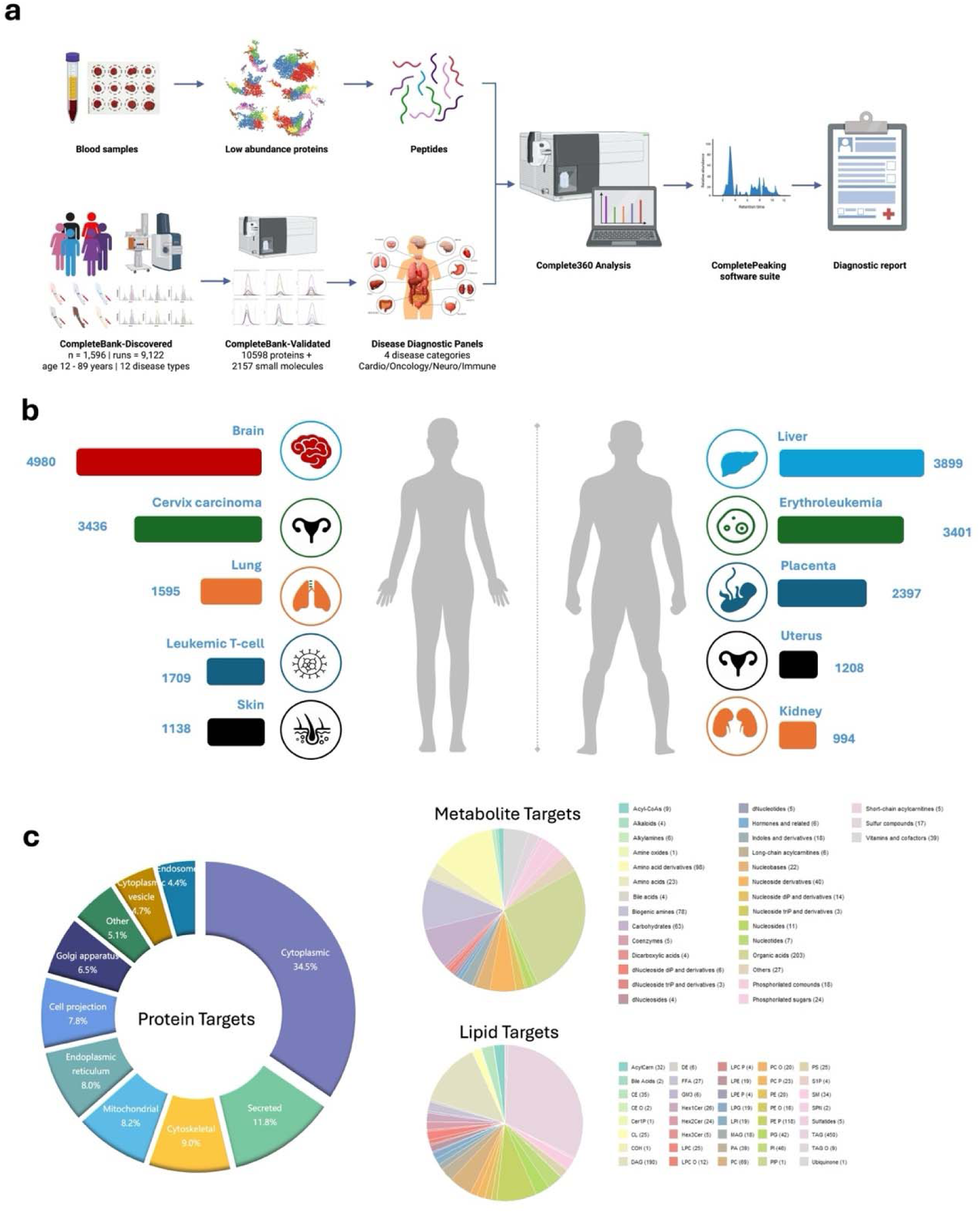
Complete360 Workflow and Target Categories. *(a)* Schematic overview of the Complete360 pipeline, starting from blood sample collection, enrichment of low-abundance proteins, enzymatic digestion to peptides, and mass spectrometry analysis. The resulting data are processed through the CompletePeaking software suite, leading to diagnostic report generation. The pipeline integrates two major reference databases—CompleteBank-Discovered and CompleteBank-Validated—and supports disease diagnostic panel development across four major categories: cardiovascular, oncology, neurological, and immune disorders. *(b)* Distribution of molecular targets across various tissues and disease-relevant organs, highlighting the breadth of proteomic coverage. Key sources include brain, liver, cervix carcinoma, lung, erythroleukemia, placenta, and kidney, among others. *(c)* Characterization of target categories: the left donut chart shows the subcellular localization of protein targets (e.g., cytoplasmic, mitochondrial, secreted); the middle and right pie charts depict the diversity of metabolite and lipid targets, respectively, classified by compound classes such as acylcarnitines, amino acids, phospholipids, and eicosanoids.

We developed a comprehensive database, termed CompleteBank, structured in two phases: CompleteBank-Discovered and CompleteBank-Validated. In the first phase, we created a discovery database encompassing 17,328 proteins and 2,927 metabolites/lipids observed in various human blood samples as described in Supplementary Table 1. This module, termed **CompleteBank-Discovered** (Figure 1a), serves as a repository for preliminary detection parameters of blood signatures, laying the groundwork for further validation. To assess the depth and uniqueness of **CompleteBank-Discovered**, we compared our protein list with the recently published Human Proteome Project (HPP) database^14^. Remarkably, we identified 536 proteins from the PE2-5 groups, marking the first-ever detection of these proteins by mass spectrometry, representing approximately 2.74% of the human proteome. This achievement is particularly significant given that these proteins were detected in blood, where protein identification is inherently more challenging. Our findings not only expand the known repertoire of detectable human proteins but also highlight the completeness and robustness of our foundational database.

The second phase focused on rigorous validation to ensure the detectability and reliability of the proteins and metabolites identified in the discovery phase. This involved a minimum of five iterative rounds of optimization for each target molecule on liquid chromatographic separation, mass spectrometric analysis, and transition selection, aimed at maximizing detection sensitivity and specificity. After each experimental round, data underwent meticulous curation. Through this process, we curated targeted mass spectrometric assays using dynamic Selected Reaction Monitoring (dSRM) corresponding to 10,598 blood proteins and 2,157 small molecules, which collectively form **CompleteBank-Validated** module (Figure 1a). This module has been developed through repeated optimizations across over 9,000 individual LC-MS/MS runs on a TOF mass spectrometer and over 1,000,000 runs on triple quadrupole (QqQ) mass spectrometers. Over 600,000 QqQ raw spectrum files were manually reviewed to ensure data quality. The **CompleteBank-Validated** module is enriched with precise detection parameters, including optimization on retention times, m/z values for MS1 and MS2 fragments, collision energies, source parameters and blood-specific noises as well as target-specific noises. It also incorporates sample preparation parameters, such as optimal sample preparation buffer systems for each individual sample type and target, and protection peptides used for each target through our proprietary MaxRec technology^12^.

To evaluate the sources from the optimized **CompleteBank-Validated** proteins, the UniProt tissue annotation database (UP_TISSUE) was used, and we found that the validated blood proteins developed in our *Complete360* platform came from all major human tissues or organs (Supplementary Table 2, Fig. 1b). Additionally, subcellular localization analysis revealed the following distribution: 34.5% cytoplasmic, 11.8% secreted, 9% cytoskeletal, 8.2% mitochondrial, 8.0% endoplasmic reticulum, 7.8% cell projection, 6.5% Golgi apparatus, 4.7% cytoplasmic vesicle, and 4.4% endosomal proteins, etc (Figure 1c); and 2,157 small molecules, where there are 762 polar metabolites from different classes (organic acids, amino acids, nucleotides, carbohydrates, etc.), covering most of the human metabolic, drug and disease pathways, and 1,395 lipid species across more than 24 (sub)classes (including aceylCarn, Bile acids, CE, CL, DAG, DE, DG, FFA, HexCer, LPC, LPE, LPG, LPI, MAG, PA, PC, PE, PG, PI, PL, PS, PIP, S1P, SM, SP, SPN, TAG, TG)^15–19^ (Figure 1c).

During the method development phase, we extensively relied on manual curation, which proved indispensable for enhancing the quality and reliability of our database. This meticulous process allowed us to fine-tune detection parameters. Over the course of approximately six years, we iteratively optimized these parameters while documenting more than 9,000 blood samples. This process enabled high sensitivity for each target detected via mass spectrometry. With these large manually curated datasets in hand, we developed an AI-based learning and peak-picking system, which we named **CompletePeaking** (Figure 1a). This system performs data analysis for each sample using comprehensive pattern-recognition algorithms to improve the identification of precise peaks for each target, with particular emphasis on determining the optimal transition(s)^20^. The manually curated spectral files served as the training set for our CompletePeaking algorithm. With CompletePeaking, the detection of all analytes has been consolidated into a suite of mass spectrometry assay clusters designed to maximize both sensitivity and reproducibility while minimizing runtime. The development of these assays incorporated several key considerations, including separation of high- and low-abundance molecules, resolution of co-eluted targets, ensure retention time reproducibility, management of co-eluting noise analytes, and minimizing Run Time, which detailed in method session.

Worthy of highlighting, as previously mentioned in management of co-eluting noise analytes, when analyzing blood samples, we consistently observed similar noise patterns across different specimens. These patterns likely originate from common high-abundance proteins or blood matrix components, such as lipids and other prevalent substances^21, 22^. These elements, while abundant, are known to limit the depth of analysis in blood proteomics and are considered detrimental to proteomic assays^23^. However, their consistent reproducibility, after being confirmed from thousands of samples, provides an opportunity to enhance data analysis automation. This consistency enables more precise peak-picking for peptide and molecule targets without the need for spiking in exogenous standards.

### Dynamic Range and Detectability of Extremely Low-Abundance Blood Proteins

Proteins in blood exhibit a vast dynamic range of concentrations, which presents a significant challenge in achieving the desired analytical depth for clinical proteomics assays. To evaluate the dynamic range of detectability provided by the *Complete360* platform, we selected a small panel of well-documented plasma proteins with known concentrations. Our findings reveal that the *Complete360* pipeline enables the detection of plasma proteins within a remarkable concentration range---from ∼10 pg/mL to ∼100 ug/mL. This dynamic range encompasses the physiological plasma concentrations of most known plasma proteins (Figure 2a).

**Figure 2.**
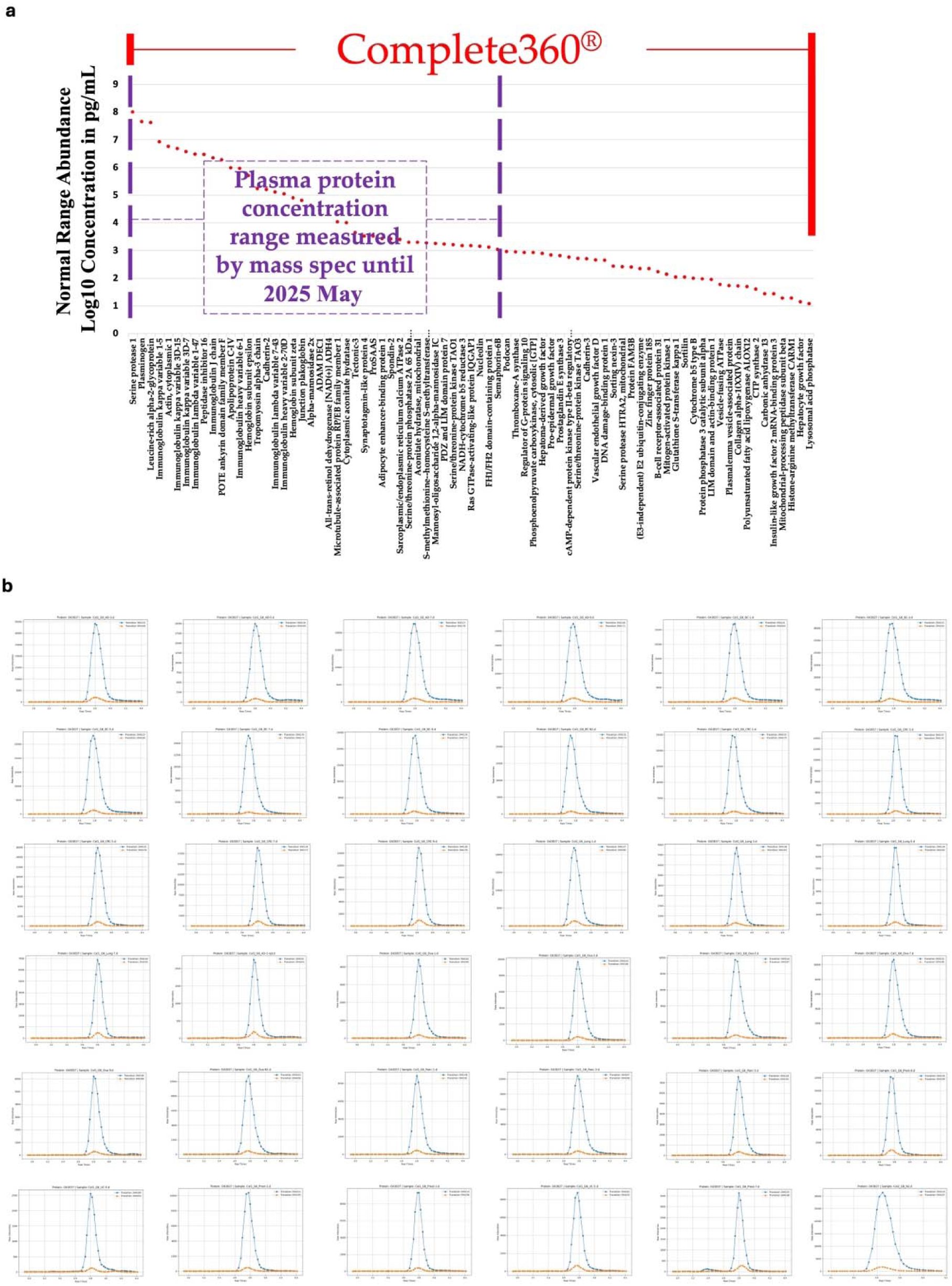
Detectability Range of Complete360®. *(a)* Dynamic range of plasma protein concentrations detectable by the Complete360 platform. The red dotted line represents the concentration distribution (Log10 scale) of a curated panel of well-characterized plasma proteins with known physiological levels. The Complete360 platform demonstrates a detection range spanning from approximately **3.5 pg/mL to over 100** μ**g/mL**, surpassing the historical detection range of mass spectrometry as of May 2025 (purple boxed area). This range captures the vast majority of physiologically relevant plasma proteins. *(b)* Representative extracted ion chromatograms for a selection of low-abundance plasma proteins detected by Complete360, including proteins reported at concentrations below 10 pg/mL. These examples highlight robust signal intensities and well-defined peaks, underscoring the platform’s ability to quantify extremely low-abundance targets with high reproducibility. Notable proteins include **Isocitrate dehydrogenase subunit beta (O43837)**, **Leukocyte cell-derived chemotaxin 1 (O75829)**, **NADH dehydrogenase flavoprotein 2 (P19404)**, **Calretinin (P22676)**, and **Methionine aminopeptidase 1 (P53582)**. These results validate the unprecedented sensitivity and analytical depth of Complete360, supporting its use in low-input and decentralized clinical applications.

One of the most challenging aspects of mass spectrometry-based detection of blood proteins is sensitivity, which depends on factors such as co-elution dynamics, contamination, and ion suppression. Detecting low-abundance protein targets has been a critical area of interest. Through repeated optimizations, we have demonstrated the ability to push the detection limits of a standard QqQ instrument to unprecedented levels at pg/mL (Figure 2a), enabling the detection of extremely low-abundance targets^6, 10–13, 24^. To further evaluate the detectability of *Complete360* for extremely low-abundance plasma proteins, we conducted assays using pooled plasma samples from healthy individuals described in previous studies^6^. The proteins detected were annotated with plasma concentration data from the Human Proteome Project (HPP) database and commercial assay providers. Notably, the lowest annotated protein detected by *Complete360* was identified at a concentration of 3.5 pg/mL (Supplementary Table 3). Furthermore, numerous proteins in the sub-10 pg/mL range exhibited robust peak profiles on the *Complete360* pipeline, underscoring its capability to detect even lower-abundance targets. For example, the protein Isocitrate dehydrogenase [NAD] subunit beta, mitochondrial (Uniprot ID: O43837) with a reported plasma concentration as 8.3pg/mL was detected in different plasma samples at high abundance, covering various disease conditions (Figure 2b). Despite its reported pg/mL level concentration, the signal intensity for this protein observed from *Complete360* was strong that it remained detectable after a 1:1000 dilution (data not shown). These findings also indicate that *Complete360* may achieve excellent detection performance even with further reduced sample input volumes. This feature has the potential to enable novel applications, such as in-home sample collection. Beyond Isocitrate dehydrogenase, several other low-abundance proteins demonstrated similarly strong intensity, including Leukocyte cell-derived chemotaxin 1 (UniprotID: O75829), NADH dehydrogenase [ubiquinone] flavoprotein 2, mitochondrial (UniprotID: P19404), Calretinin (UniprotID: P22676), and Methionine aminopeptidase 1 (UniprotID: P53582), etc.

### High Reproducibility

Reproducibility is a critical requirement for clinical diagnostics, as it ensures precision and reliability in assay results. With clinical applications as our focus, we systematically evaluated the reproducibility of the *Complete360* assay using two complementary approaches:

We evaluated the detection and quantification of 36 proteins across 12 replicates. These proteins span a wide dynamic range of documented plasma concentrations, from 19 pg/mL to 25 ug/mL, covering over 10 orders of magnitude (Table 1). By including proteins at varying concentrations in plasma, we carefully assessed reproducibility across a wide dynamic range, highlighting the robustness and consistency of the *Complete360* platform. Remarkably, the results demonstrated high reproducibility, with an average coefficient of variation (CV) in quantification of only 3.92% ranging from 1.4% to 7.0% across all proteins on the panel (Table 1). Representative raw spectra of key proteins illustrate this exceptional reproducibility (Supplementary Figure 1). These findings demonstrate that the *Complete360* assay is highly reproducible, meeting the stringent requirements necessary for clinical applications.

To assess a broader range of proteins, we conducted *Complete360* assays targeting 9,977 proteins across five replicates (Supplementary Table 4). The median CV for the entire panel was 11.97%. When focusing on proteins with CVs below 25%, 7,833 proteins were consistently detected across all replicates, with a median CV of 8.73%. Notably, 4,361 proteins exhibited CVs below 10%, with a median CV of 4.77%. This subset of highly reproducible proteins demonstrates significant potential for direct translation into clinical applications once a clinical relevance to a disease is validated.

It is noteworthy that the reproducibility of protein abundance measurements and signal intensity detected by the *Complete360* platform demonstrates a strong correlation (Figure 3). As the biological concentration of proteins in plasma increases, the reproducibility of detection improves accordingly. Moreover, the *Complete360* platform reliably quantifies protein targets across a dynamic range exceeding eight orders of magnitude. The correlation between QqQ intensity and reproducibility is remarkably high, with an R² value of 0.51. It is also important to highlight that only about 4,500 proteins have documented blood concentration data to date^25^. Interestingly, most of the proteins included in the *Complete360* full panel are not yet documented for their blood concentration levels. Future efforts will focus on systematically establishing these concentration profiles to further enhance the clinical and diagnostic utility of our platform, including achieving reliable absolute quantifications.

**Figure 3.**
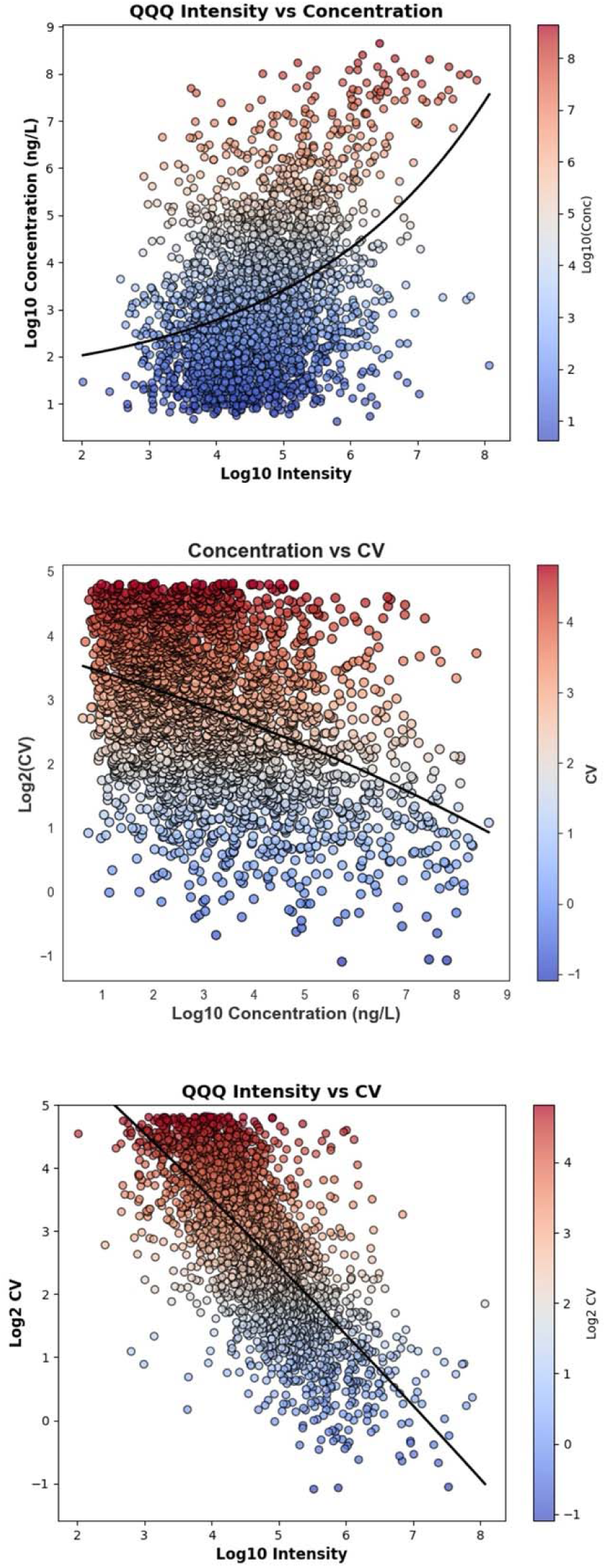
CV of Quantification Across Different Mass Spectrometry Intensities and Protein Concentrations. This figure shows how quantification reproducibility (coefficient of variation, CV) relates to mass spectrometry intensity and protein concentration using the Complete360 platform. *(Top)* QqQ intensity correlates positively with known plasma protein concentrations, spanning a wide dynamic range. *(Middle)* Higher protein concentrations are associated with lower CVs, indicating better quantification reproducibility. *(Bottom)* A strong inverse correlation (R² = 0.51) is observed between QqQ intensity and CV, confirming that higher signal intensity leads to more reliable protein measurement.

Next, we compared the reproducibility and detectability of *Complete360* to conventional DIA-based proteomic profiling. Using timsTOF HT to analyze the same sample, we conducted three technical replicates, resulting in the identification of 7,697 proteins using DIA-MS. (Supplementary Table 5). Among these, 3,944 proteins were consistently detected across all three replicates, 2,737 proteins were observed in two out of three replicates, and 1,016 proteins were identified in only one replicate. For proteins detected in two or more replicates, the median CV for quantification was calculated to be 15.23%, indicating a high-quality profiling assay. These results demonstrate that *Complete360*, with its optimized parameters, provides substantially enhanced reproducibility and quantification consistency compared to conventional DIA-based approaches.

Our findings demonstrated the reproducibility of the *Complete360* platform across both small-and large-scale protein panels, covering a wide dynamic range of protein concentrations. The ability to consistently achieve results with CVs well within acceptable thresholds highlights *Complete360* as a reliable and practical solution for clinical applications. Its enhanced reproducibility and sensitivity, offering improved reliability and confidence in detection and quantification plasma proteins, making *Complete360* a valuable tool for mass spectrometry-based proteomics, particularly in clinical diagnostics where precision and consistency are essential.

### Disease-Associated Molecular Changes Revealed by Complete360

Using the *Complete360* platform, we applied *Complete360* assays on plasma samples, representing patients diagnosed with breast cancer and non-cancerous control (Supplementary Table S6).

The comparison between the plasma proteins from breast cancer and non-tumor samples using the *Complete360* assays could lead the discovery of specific plasma protein changes for breast cancer. The significantly altered plasma proteins between tumor and non-tumor samples could be potentially useful for the diagnosis of breast cancer. In the discovery plasma proteomic data set using Evelyn MS instrument, the principal component analysis (PCA) of breast cancer (n=5) and non-cancerous samples (n=7) illustrated a formation of distinct clusters of the cancer and non-cancerous samples (Figure 4A). All cancer samples were differentiated from the non-cancerous samples. From the differential analyses of breast cancer and non-cancerous control plasma samples, we identified 331 up-regulated and 284 down-regulated plasma proteins (Supplementary Table S6). To verify the identified plasma protein changes, we analyzed additional plasma samples from 5 independent breast cancer patients and 9 non-cancerous plasma samples by *Complete360* assays with a orthogonal MS instrument, Amelia (Supplementary Table S6). The PCA analysis showed the formation of distinct clusters of the cancer and non-cancerous samples (Figure 4B). We found that the quantitative data from the independent patient plasma with orthogonal MS instrument is consistent with the discovery analysis with correlation of 0.54 (Figure 4C). Using the verification breast cancer data, 105 of the 331 elevated plasma proteins identified from discovery data were verified by the independent analyses of the additional samples using *Complete360* assays with a different MS instrument (Supplementary Table S6).

**Figure 4.**
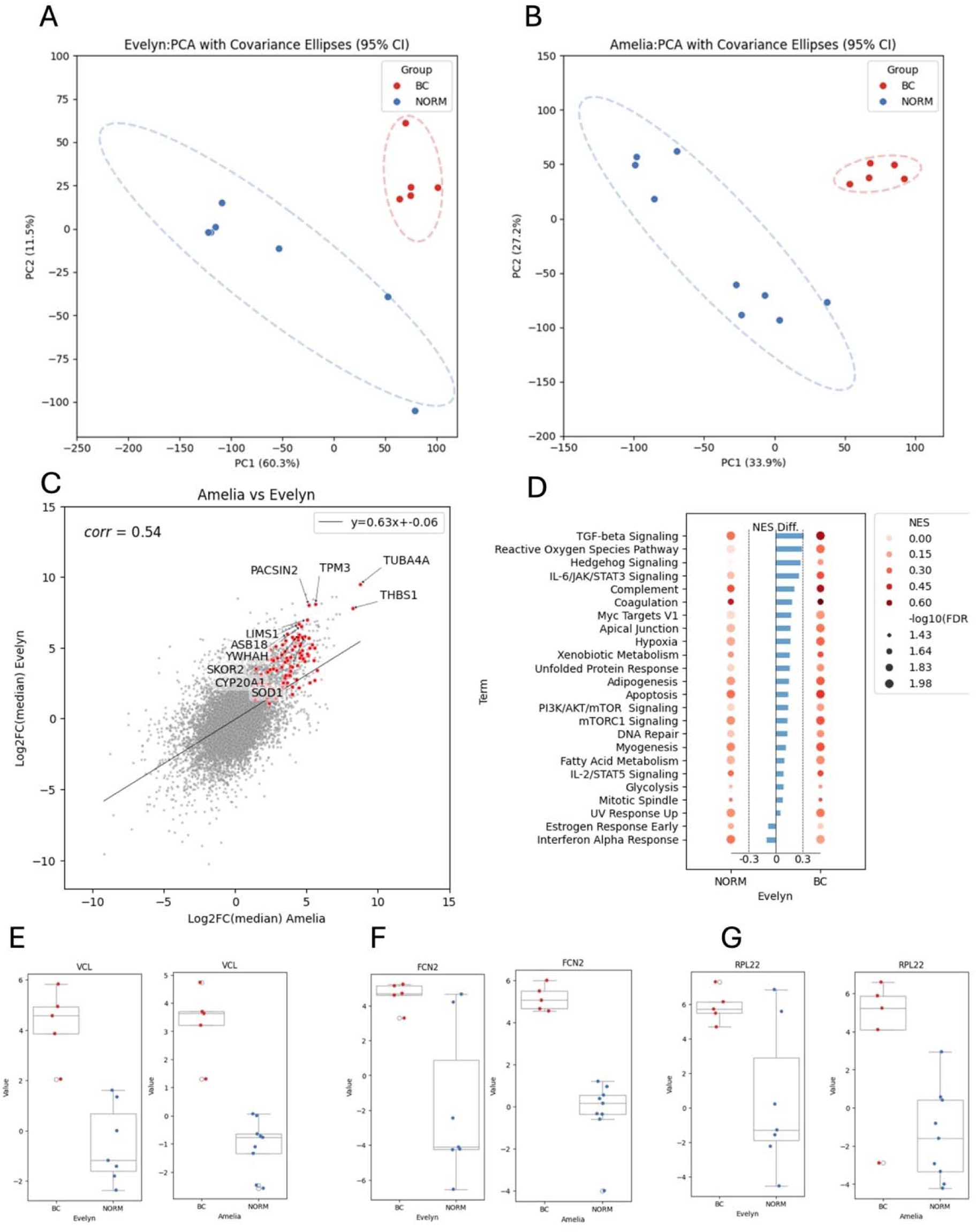
Disease-Associated Molecular Changes Revealed by Complete360®. *(A–B)* Principal component analysis (PCA) of plasma proteomic profiles from breast cancer (BC) and non-cancer (NORM) samples in two independent cohorts— Evelyn *(A)* and Amelia *(B)*—demonstrates clear separation between disease and control groups, indicating robust disease-associated proteomic signatures. *(C)* Correlation plot comparing log2 fold changes of proteins between the Amelia and Evelyn cohorts. Red-highlighted proteins denote those significantly upregulated in breast cancer samples across both cohorts. Key tumor-associated proteins are labeled, including **THBS1**, **TUBA4A**, **TPM3**, **PACSIN2**, and **SOD1**. *(D)* Single-sample gene set enrichment analysis (ssGSEA) of the Evelyn cohort, using all quantified plasma proteins and the 50 Hallmark gene sets from the MSigDB Hallmark 2020 database. *(E–G)* Boxplots showing expression levels of three representative upregulated proteins in breast cancer plasma samples compared to non-cancer controls, such as (E) *Vinculin (VCL)*, (F) *Ficolin-2 (FCN2)*, and (G) *Ribosomal protein L22 (RPL22)*.

To investigate the classes of plasma proteins that were specifically regulated in breast cancer, the Hallmark pathways were performed on the significantly positive regulated plasma proteins (Supplementary Table S6, Figure 4D) and revealed that TGF-beta signaling, reactive oxegen species pathway, IL-6/JAK/STAT3 signaling, complement, Myc targets are top overrepresented pathways for plasma proteins elevatged in breast cancer, while estrogen responses early, interferon alpha response were overrepresented in significantly down-regulated proteins for breast cancer.

The comparison between the plasma proteins from breast cancer and non-cancerous samples could lead the discovery of specific protein changes for breast cancer and serve as protein markers for breast cancer detection using blood samples. Among the upregulated plasma proteins in breast cancer plasma, several key proteins linked to tumor progression and metastasis were identified. Notable proteins include **vinculin (VCL, Figure 4E))**, involved in cell migration and metastasis ^26, 27^. **Ficolin-2 (FCN2, Figure 4F)**, an immune system protein, plays a role in immune surveillance, and **Large ribosomal subunit protein eL22 (RPL22, Figure 4G)**, frequently deregulated in cancers, suggests its role in tumor biology ^28, 29^.

### Simultaneous Quantification of Plasma Proteins, Metabolites, and Lipids for Enhanced Diagnostic Precision

To achieve comprehensive diagnostics, we have integrated metabolomics and lipidomics analysis within the *Complete360* platform, enabling the simultaneous detection and quantification of proteins, metabolites, and lipids from the same biological sample through the same platform. This streamlined multi-omics approach maximizes sample utilization, enhances diagnostic accuracy while maintaining cost efficiency. By consolidating all assays onto a unified platform, our method facilitates seamless clinical implementation, supporting broader adoption in clinical and translational research.

Our current *Complete360-MyMeta* assay employs a targeted method capable of detecting 762 metabolites and 1,395 lipids, with systematic optimization across multiple refinement cycles to enhance sensitivity and reproducibility. The metabolomics assays cover both polar metabolites and lipids, with data undergoing separate median normalization for metabolites (Supplementary Table 7) and lipids (Supplementary Table 8).

For each disease, our integrated approach has yielded highly informative results, revealing matched metabolic and proteomic signatures (Supplementary Table 9). Notably, pathway analysis shows strong concordance between proteomic and metabolomic data, with an average of 69% of the top 10 pathways concerning metabolomics identified from each omics perspective overlapping. This alignment underscores the robustness and biological relevance of our integrated *Complete360* platform in disease characterization.

To fully leverage the capabilities of the multi-omics *Complete360* platform, we integrated proteins, metabolites, and lipids as key features and performed a *t*-test to compare each disease against all other conditions. Molecules were ranked in ascending order based on p-values, and the top 1,000 features were consistently selected for model construction. We then generated ROC curves using only the top 1,000 proteins for each disease and compared them to ROC curves generated when the top 1,000 features—comprising proteins, polar metabolites, and lipids—were collectively incorporated (Figure 4b). To ensure consistency, the total feature count was always maintained at 1,000, allowing the model to determine the optimal composition of proteins, metabolites, and lipids for each disease-specific diagnostic panel. On average, 794 proteins were selected, while the mean feature count for metabolites and lipids was 90 and 117, respectively (Supplementary Table 10). Notably, we observed an increase in AUC values for most diseases when multi-omics features were incorporated into the diagnostic model, underscoring the advantage of integrating multiple molecular layers (Figure 4b).

This enhancement highlights the unique value of the *Complete360* platform, which enables the simultaneous and cost-effective analysis of multi-omics analytes on a single instrument, improving both diagnostic accuracy and efficiency. We acknowledge that the current study is based on a limited sample size, and while the observed ROC curves provide valuable insights, they may not fully capture the diagnostic potential of our approach. Future studies with larger sample cohorts and deeper data analysis approaches will be conducted to further validate these findings, as this work primarily serves as a proof-of-concept demonstration of the *Complete360* platform.

### Plasma Proteome Variation and Its Genetic Determinants Revealed by Complete360

Using *Complete360* methods, we have conducted an ultra-deep plasma proteomics analysis to investigate the correlation between plasma protein levels and human age, gender, and BMI. Our findings align closely with those reported by Mann *et al.*, demonstrating that a significant proportion of the plasma proteome varies systematically with these demographic and physiological factors^30^. Notably, we identified age-, gender- or BMI-associated shifts in proteins involved in inflammation, extracellular matrix remodeling, lipid metabolism, and coagulation cascades, reflecting the dynamic changes in systemic physiology (Figure 5).

**Figure 5.**
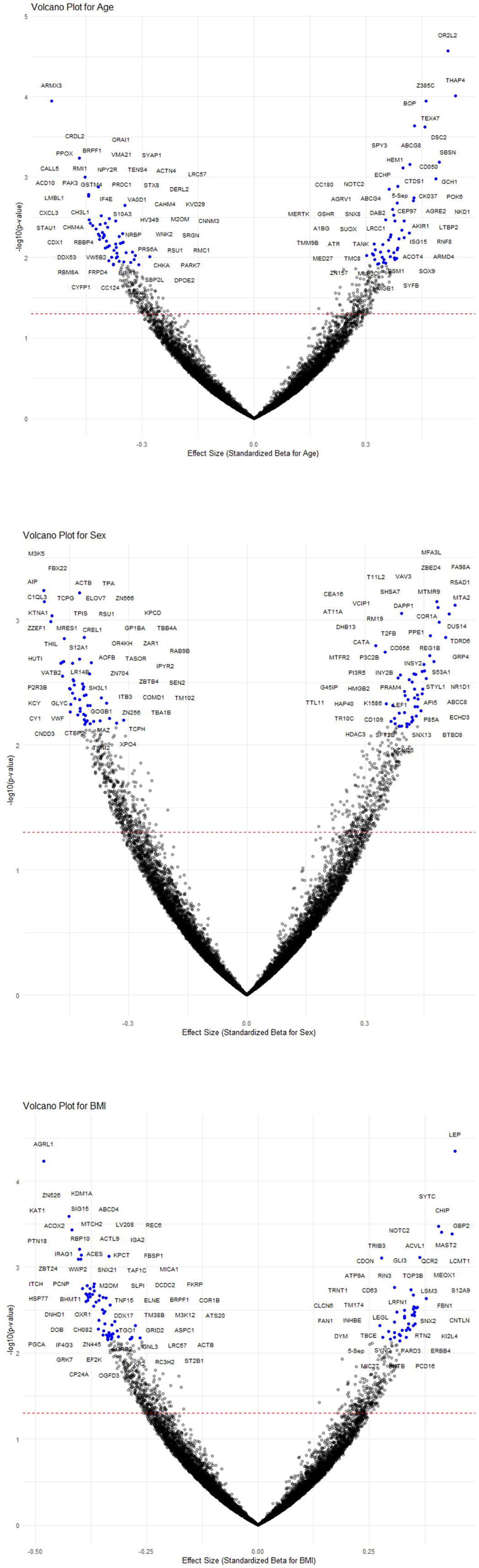
Complete360-Based Quantitative QTL (pQTL) Analysis for Age, Gender, and BMI. Volcano plots showing the association between plasma protein expression levels and three key demographic and physiological traits—age, sex, and body mass index (BMI)—as determined using the Complete360 ultra-deep proteomics pipeline. *(Top)* Volcano plot for age-related proteomic variation. Multiple proteins involved in inflammation, extracellular matrix remodeling, and cellular senescence display significant age-associated shifts, indicating the utility of Complete360 in capturing dynamic physiological aging markers. *(Middle)* Volcano plot for sex-based differences in plasma proteomes. Sex-specific regulation is observed in proteins related to hormonal signaling, immune response, and lipid metabolism. *(Bottom)* Volcano plot for BMI-associated proteomic changes. Proteins such as **TRIB3**, **INHBE**, and **ERBB4** show strong BMI-correlated expression profiles, while **LEP (leptin)** emerges as a key biomarker for adiposity and metabolic regulation. These findings reveal broad and systematic proteomic variation linked to age, sex, and BMI, highlighting the precision and reproducibility of the Complete360 platform. The ability to detect physiologically relevant protein signatures supports the platform’s utility for biomarker discovery, personalized health monitoring, and metabolic disease risk stratification.

*Complete360*’s high-sensitivity protein detection enabled the identification of BMI-associated proteomic signatures, uncovering key proteins involved in metabolic regulation, inflammatory response, and lipid transport (Figure 5). Notably, proteins such as TRIB3 (a regulator of obesity and insulin resistance), INHBE (a determinant of fat distribution), and ERBB4 (which modulates brain-regulated energy expenditure) exhibited distinct expression profiles across BMI categories. Among these, LEP (leptin) emerges as a particularly significant contributor to BMI, reinforcing its well-documented role in weight regulation^31^. These findings reveal a robust molecular signature of metabolic health, providing a valuable framework for biomarker discovery and disease risk stratification. The strong correlation between these plasma proteins and BMI underscores the predictive power of *Complete360* in distinguishing metabolic states. This highlights the potential of plasma proteomics not only as a biological clock for metabolic health but also as an innovative tool for early disease detection and personalized health monitoring.

Furthermore, *Complete360* facilitates genetic-proteomic association studies (pQTL analysis) to determine the genetic influences on plasma protein levels. Our initial findings suggest that genetic variants contribute significantly to the observed protein-level variance, with some proteins showing strong cis- and trans-regulatory effects. The integration of *Complete360* with genome-wide association studies (GWAS) is expected to further uncover causal relationships between genetic factors, proteomic alterations, and disease predisposition.

With its ability to quantify thousands of plasma proteins at high specificity, capture proteomic variability with minimal technical noise, and support predictive modeling of age and BMI, *Complete360* is at the forefront of precision medicine and multi-omics biomarker research. These insights will be instrumental in improving disease risk assessment, enhancing therapeutic targeting, and advancing our understanding of human health at the molecular level.

### *Complete360* Assay for Dried Blood Spot Samples

Dried blood spot (DBS) sampling is a widely adopted method for at-home sample collection due to its convenience, ease of storage, and cost-effective transportation via standard mailing services. However, proteomics assays face substantial challenges when applied to DBS samples. Affinity-based proteomics methods often suffer from loss of protein epitope integrity and higher-order structural degradation during prolonged room-temperature storage. Similarly, mass spectrometry-based approaches are hindered by an overwhelming release of peptides from red blood cells and other cellular components, compromising the depth and specificity of plasma proteome analysis derived from whole blood DBS samples. The *Complete360* platform overcomes these limitations for DBS analysis. Unlike affinity-based methods that depend on intact epitopes, *Complete360* utilizes a proprietary approach optimized for detecting targets that are inherently resistant to proteolysis and chemical modifications. These targets have been carefully selected and refined to enhance assay performance. Proteins released into the sample due to prolonged storage at room temperature do not interfere with the detection of desired targets, allowing the assay to maintain high sensitivity and specificity.

To assess the assay’s performance under varied storage conditions, DBS samples were collected and stored at ambient temperature for 1 to 12 days in standard mailing envelopes, simulating routine transport scenarios (Figure 6a). The samples were processed using *Complete360* platform (Materials and Methods, Supplementary Figure 2) and analyzed for proteins and small molecules using the *Complete360* pipeline.

**Figure 6.**
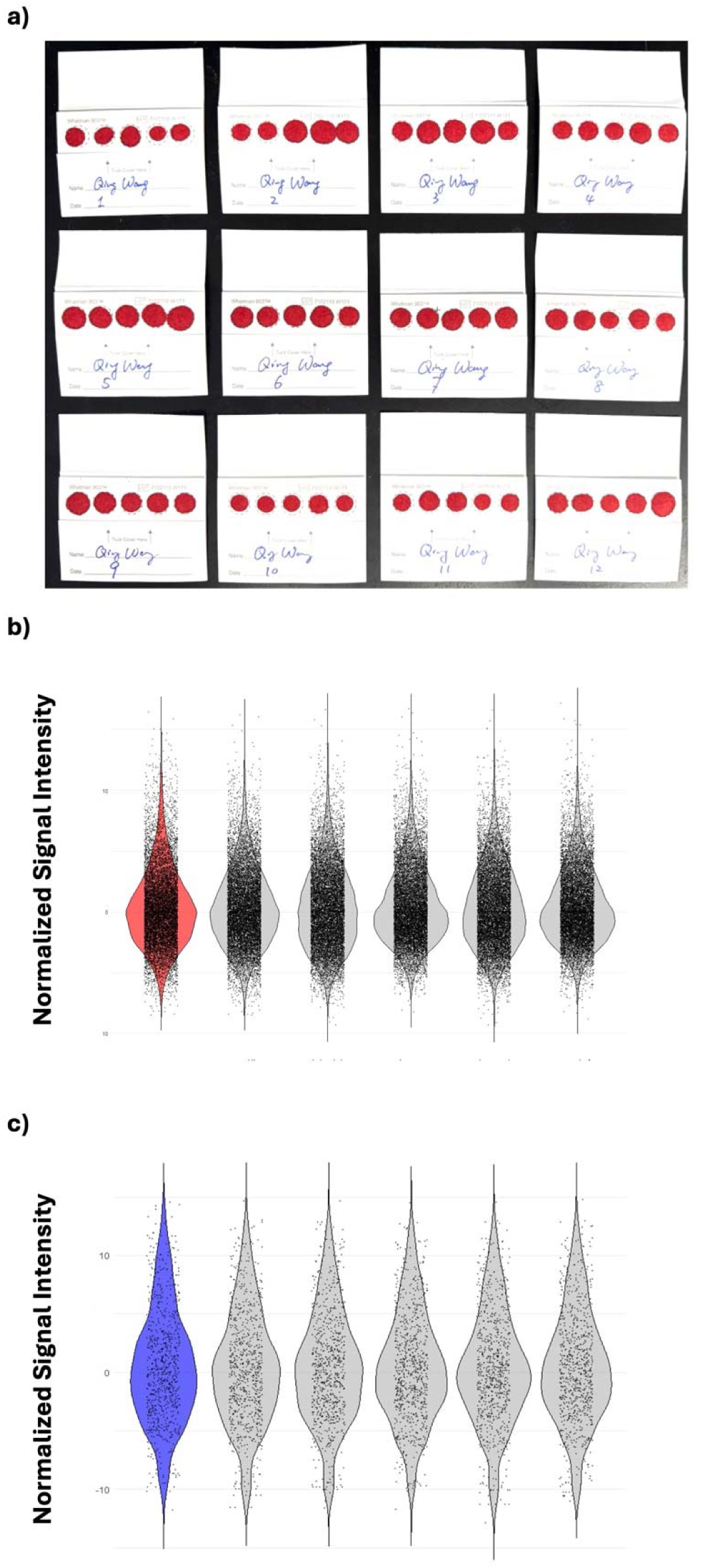
Analysis of Dried Blood Spot (DBS) Samples Using the Complete360 Pipeline. *(a)* Dried blood spot (DBS) samples were collected from the same individual and stored at room temperature for varying durations, as indicated on the collection card, prior to analysis. *(b)* Comparison of protein biomarker intensities between matched plasma and DBS samples demonstrates high consistency, validating the robustness of the Complete360 pipeline in protein quantification from DBS. *(c)* Metabolite and lipid biomarker intensities from matched plasma and DBS samples also show strong correlation, supporting the platform’s ability to perform integrated proteomic and metabolomic profiling from minimal, remote-collected samples.

Through *Complete360*, we systematically analyzed over 10,000 proteins from DBS samples. Approximately 2–3% of proteins exhibited consistent temporal changes, with 340 proteins showing a steady increase and 279 proteins showing a continuous decrease over storage periods of 1, 3, 5, 7, and 9 days. Additionally, 46.96% of proteins maintained a coefficient of variation (CV) below 25% across all DBS samples collected from the same individual, demonstrating the pipeline’s stability and resistance to long-term exposure to air and room temperature (Supplementary Table 11). Notably, plasma protein profiles from DBS samples closely matched those from conventionally collected plasma samples from the same individual (Figure 6b-c), supporting the reliability of *Complete360* for proteomic analysis of DBS samples. However, further studies are needed to evaluate the diagnostic performance of using diseased DBS samples, as this was beyond the scope of the current study.

Based on these data, we developed a normalization reference dataset encompassing all *Complete360*-detected proteins over a 12-day period from the same individual (Supplementary Table 11). This dataset enables future analyses to account for time-dependent variations in protein abundance, enhancing the accuracy of disease biomarker quantification from DBS samples. By integrating the mailing timestamp of DBS samples, this approach allows for precise adjustments to compensate for protein changes occurring during shipment and storage to best reflect the original clinical status of the patient when the DBS sample was collected.

Furthermore, a discovery-mode analysis using the timsTOF HT platform identified 5,781 proteins from DBS samples, significantly expanding the depth of coverage compared to conventional plasma proteomics (Supplementary Table 12). Notably, 133 proteins were uniquely detected in DBS samples and had not been previously observed in freshly frozen plasma and is outside of our CompleteBank database of 17,328 plasma proteins.

DBS is an inherently robust sample type for metabolomics and lipid assays. Using *Complete360*, we identified a comprehensive set of 1,395 metabolites and 762 lipids from DBS samples, demonstrating the feasibility of high-throughput multi-omics analysis. Notably, the levels of these small molecules remained remarkably stable across multiple days (Supplementary Table 13), underscoring the reliability of DBS for longitudinal studies. These findings highlight the potential of DBS-based *Complete360* analysis for clinical applications in disease detection and monitoring, with its utility strongly supported by the stability of the detected analytes in this dataset.

These findings underscore the exceptional versatility and robustness of the *Complete360* platform in accommodating diverse sample collection and storage conditions, including long-distance transport and room-temperature storage of DBS samples. This capability has the potential to redefine clinical proteomics by enabling comprehensive multi-omics analyses from easily accessible and transportable biological specimens. The ability to derive deep proteomic and metabolic insights from DBS samples opens new avenues for remote patient monitoring, large-scale epidemiological studies, and global healthcare applications, ensuring the generation of high-quality multi-omics data regardless of logistical constraints.

## Discussion

*Complete360* is a highly targeted and comprehensive detection platform, capable of quantifying close to 13,000 molecules from blood samples, delivers unmatched sensitivity and reproducibility, exceeding the capabilities of traditional profiling methods commonly used in academic and clinical settings. Underpinned by the comprehensive CompleteBank databases and CompletePeaking algorithms, *Complete360* establishes itself as a potential transformative tool for basic research and clinical diagnostics, offering advancements in biomarker discovery, disease pathway analysis, and personalized medicine. It is designed to bridge the gap between multi-omics research and real-world clinical applications, enabling newly identified molecular changes to be seamlessly translated into clinical use on the same platform.

Proteomics assays generally follow two main approaches: mass spectrometry-based methods and affinity-based detection techniques. Mass spectrometry relies on advancements in instrumentation, sample preparation, and data analysis algorithms, while affinity-based methods employ antibodies or aptamers to facilitate assays such as ELISA or its variations, like proximity extension assays (PEA). Although affinity-based methods have been widely applied to clinical specimens, their limitations are apparent. They rely heavily on the quality of the binding reagents, which can lead to inconsistencies due to variations in the manufacturing of antibodies or aptamers^32–34^. Even with high-quality binding reagents, the detection may be hindered by the limited accessibility of epitopes; many proteins in blood are modified by different protein modifications or form complexes by binding to other molecules, obscuring their binding sites^35^. Furthermore, many proteins that serve as valuable biomarkers and are involved in rapid physiological responses have relatively short half-lives therefore hindering their detection by affinity-based methods^36, 37^. For example, insulin and glucagon, both critical for glucose regulation, have half-lives of about 5 to 10 minutes, while cytokines like interleukin-6 can range from minutes to a few hours. Although these short-lived proteins are essential disease biomarkers, accurately detecting them using affinity-based methods is challenging. This is due to epitope masking through binding to other proteins, or rapid epitope damage and degradation caused by protease digestion. These factors often result in compressed fold-change data in affinity-based methods, making it difficult to differentiate between disease and healthy individuals. As a result, the sensitivity and specificity required for effective diagnostics are significantly compromised.

Mass spectrometry-based proteomics offers superior specificity and resolution, making it well-suited for distinguishing disease from control samples. However, these methods also face challenges related to sensitivity and reproducibility. Most proteomics assays employ profiling techniques using orbitrap or time-of-flight mass analyzers, and these platforms often fall short of the reproducibility standards required for clinical applications, where a coefficient of variation (CV) below 10% is essential. While triple quadrupole mass spectrometers are widely used in clinical laboratories to detect disease-associated small molecules, they require extensive optimization of detection parameters, including sample preparation strategies. Despite efforts to establish standardized detection protocols using synthetic peptides, these parameters often remain theoretical and may not fully account for the variability of real-world clinical samples.

Given these challenges, there is an urgent need for a robust and reproducible proteomics platform capable of detecting a broad spectrum of clinically relevant proteins and metabolites from blood samples with high accuracy and reliability. This is the driving force behind the development of *Complete360*. The platform is designed to provide a finely tuned, clinically viable system for comprehensive proteomic and metabolic analysis in blood, ensuring the reproducibility and sensitivity required for clinical applications. Through years of refinement, we have optimized sample preparation workflows, established precise detection parameters, and developed a sophisticated data analysis pipeline. Validated through the analysis of a good amount of body fluid samples, *Complete360* represents a major advancement in proteomics research and its translation into clinical practice. Looking ahead, we aim to extend the application of *Complete360* beyond basic research to direct clinical diagnostics. Our goal is to implement this platform across multiple countries, facilitating improved disease detection and better patient outcomes. By bridging the gap between proteomics research and clinical application, *Complete360* has the potential to redefine the future of precision medicine.

## Materials and Methods

### Ethical statement

This study was approved by the Institutional Review Boards for Human Research at Complete Omics Inc. (Baltimore, MD, USA) and BioIVT (Westbury, NY, USA), and complied with the Health Insurance Portability and Accountability Act. We collected comprehensive clinical and demographic information, medical history, comorbidity information, and vaccination schedules for all patients and participants. All participants provided informed consent to participate.

### Chemicals and Reagents

For blood proteomics: Water Optima LC/MS grade; Acetonitrile Optima LC/MS grade; Methanol LC/MS grade; Ammonium Bicarbonate (ABC) 1M; Tris buffered saline (Sigma), Formic Acid 98%-100%; Sodium dodecyl sulfate (SDS); Tris-(2-Carboxyethyl)phosphine-HCl (TCEP); 2-Chloroacetamide (CAA) ≥98%; Triethylammonium bicarbonate (TEAB) 1.0 M; Triethylamine; Phosphoric Acid; Promega sequencing grade Trypsin; Whatman 903TM blood collection kit, Minute^TM^ albumin depletion kit.

For small molecules: Ammonium formate, Ammonium acetate, Ammonium hydroxide solution: Sigma-Aldrich; Methanol (LC), water (LC/MS Grade), acetonitrile, and 2-propanol (LC/MS grade, LiChrosolv): Fisher Sci.

### Patients and samples

Plasma samples used in this study were obtained from BioIVT (Westbury, NY, USA), along with comprehensive clinical information (Supplementary Table 14). All samples were collected in accordance with institutional ethical guidelines and were de-identified to ensure patient confidentiality. Plasma was collected using purple-top tubes containing EDTA as an anticoagulant. Upon collection, samples were processed promptly by centrifugation to separate plasma, aliquoted, and stored at −80°C until further use to minimize freeze-thaw cycles and maintain proteomic and metabolic stability.

### Plasma sample preparation and analysis methods

Plasma samples were processed using the *Complete360*-MyProt pipeline, incorporating our proprietary Chemical-Biological Plasma Protein Preparation procedure. This workflow starts from two key steps to remove high abundance proteins and collect clinically and biologically more meaningful low-abundance proteins:

1. Chemical Procedure: Major plasma proteins were precipitated using a set of in-house-prepared protein removal reagents.
2. Biological Procedure: The remaining high-abundance and median-abundance plasma proteins were depleted using a proprietary antibody-conjugated resin, targeting a combination of proteins that are most frequently detected by mass spectrometry and reported in the database of peptide atlas. Such depletion procedure has been observed to be more reproducible for protein quantifications compared to that of nanoparticle-based plasma low-abundance protein enrichment methods (data not shown). Protein depletion methods for other body fluid can be established the same way. The depletion resin is tested for durability, demonstrating consistent performance for over 200 uses with optimized buffers and procedures (data not shown) to ensure an ultra-low cost for plasma protein extraction.

After the removal of high- and median-abundance proteins through this chemical-biological procedure, the plasma proteins remaining in the supernatant were processed into peptides using the *Complete360* sample digestion kits and reagents. Briefly, plasma proteins were denatured using SDS and digested with an optimized trypsin digestion protocol. Following digestion, the resulting peptide samples were fractionated using an offline HPLC system operating in both low-pH and high-pH modes. This dual-mode approach ensures highly reproducible chromatographic profiles. The procedures were extensively optimized for human plasma samples, with key metrics such as protein identification, detected abundance, and mis-cleavage rates carefully monitored to ensure reproducibility and sensitivity of mass spectrometry analysis.

For proteomics analysis using dried blood spot (DBS) samples, three 12 mm disks were pooled and incubated in Tris-buffered saline (TBS) containing 0.05% NP-40 at 37 °C for 30 minutes with agitation. The supernatant was then combined with an equal volume of the Minute™ Albumin Depletion Kit reagent to remove albumin and hemoglobin; this depletion step was repeated twice. The resulting precipitate was solubilized in TBS and subjected to digestion using *Complete360* sample digestion kits and reagents as described above.

Peptide digests were then subjected to basic reversed-phase chromatography using an Agilent 1260 liquid chromatography system, following the methodology outlined reported previously^38^. Separation was performed on an in-house packed C18 column employing a gradient of acetonitrile in 10 mM triethylammonium bicarbonate (TEAB). The gradient conditions were as follows: 5% to 28% solvent B over 75 minutes, increased to 42% over the next 8 minutes, and then to 98% over the subsequent 3 minutes, at a flow rate of 1 mL/min. Fractions were collected every minute and were then concentrated to dryness using a SpeedVac equipped with a chilled vacuum trap. The dried peptides were stored at −80 °C until further analysis.

DIA-MS Analysis: Mass spectrometric discovery analyses were conducted using a timsTOF HT mass spectrometer coupled to a nanoElute 2 liquid chromatography system via a CaptiveSpray™ ion source, configured in a two-column setup comprising a 5 mm Thermo trap cartridge and a PepSep Max Ten series analytical column (10 cm × 150 µm i.d., 1.5 µm particle size).

DDA-PASEF Analysis: To assess the quality of trypsin digestion, including the evaluation of missed cleavages and potential artifacts, data-dependent acquisition parallel accumulation–serial fragmentation (DDA-PASEF) analyses were performed. This approach facilitated the identification of peptides and proteins, ensuring the integrity of the digestion process.

DIA-PASEF Analysis: For comprehensive proteomic profiling, data-independent acquisition PASEF (DIA-PASEF) analyses were executed. The acquisition method was optimized to minimize missing data and to cover ion mobility ranges with high-density precursor sampling. The method consisted of eight cycles, each comprising 29 ion mobility (IM) windows. An initial MS1 scan was followed by eight DIA-PASEF cycles, covering an m/z range of 375–1100 and an inverse reduced mobility (1/K) range of 0.65–1.45 V·s/cm². The resulting DIA-MS data were processed using DIA-NN (version 1.8.2) employing a predicted human protein spectral library containing 20,480 entries. Both DDA- and DIA-PASEF datasets were analyzed using DIA-NN and FragPipe software to ensure comprehensive identification and quantification of peptides and proteins^39, 40^.

*Complete360*-MyProt Analysis: Targeted detection and quantification for plasma proteins was performed on an Agilent 6495 QqQ Mass Spectrometer using dynamic Selected Reaction Monitoring (dSRM) with an Agilent Jet Stream ion source. Chromatographic separation was achieved using an in-house packed C18 column (1.7 µm, 2.1 mm × 30 mm). The gradient of solvent B (acetonitrile with 0.1% formic acid) was programmed as follows: 12% to 42% over 5.4 minutes, followed by an increase to 98% in the next minute, at a flow rate of 150 µL/min. A set of targeted assays were created through CompleteBank-Discovered and CompleteBank-Validated process to therefore detect and quantify over 10,000 plasma proteins in the same assay.

### Sample Preparation and Analysis Methods for Small Molecules

Plasma samples were processed using the *Complete360*-MyMeta pipeline using a modified MTBE/MeOH/H2O extraction protocol to isolate lipids and polar metabolites from 40ul plasma sample. First, 300 µL of cold methanol was added to the plasma aliquot and vortexed for 10 seconds. Following this, 1 mL of methyl tert-butyl ether (MTBE) was added, and the mixture was vortexed again for 10 seconds before being incubated on a shaker at room temperature for 60 minutes. After incubation, 250 µL of water was added to induce phase separation, followed by a 10-minute incubation at room temperature with occasional vortexing. The samples were then centrifuged at 15,000 g for 10 minutes to separate the phases. The upper organic lipid phase (approximately 900 µL) was collected into a clean 2 mL glass vial, while the lower aqueous metabolite phase (320–350 µL) was also collected. The organic phase was dried under nitrogen gas for 20-30 minutes and reconstituted in 300 µL of 1-butanol/methanol (1:1) containing 10 mM ammonium formate. The aqueous phase was dried under nitrogen and reconstituted in 150 µL of 50% acetonitrile (ACN).

#### *Complete360*-MyMeta Analysis

For MyMeta analysis, metabolites were separated on an Agilent 1290 LC system using two HILIC-LC methods. The first method, HILIC-01, employed an Acquity BEH-Amide column (1.7 µm, 2.1 × 150 mm) at a column temperature of 40 °C and an autosampler temperature of 8 °C. The injection volume was 5 µL, with mobile phase A consisting of 95% water + 20 mM ammonium acetate (pH 9.4), and mobile phase B being 98% acetonitrile. The flow rate was set at 0.15 mL/min with a gradient program running as follows: 0 minutes, 90% B; 2 minutes, 90% B; 3 minutes, 75% B; 7 minutes, 75% B; 8 minutes, 70% B; 9 minutes, 70% B; 10 minutes, 50% B; 12 minutes, 50% B; 13 minutes, 25% B; 14 minutes, 25% B; 16 minutes, 0% B; 20 minutes, 0% B; 21 minutes, 90% B; and 25 minutes, 90% B, with a 2-minute post-run period. The second method, HILIC-02, used an Atlantis Premier BEH Z-HILIC column (1.7 µm, 2.1 × 100 mm) at a column temperature of 30 °C and an autosampler temperature of 8 °C. The injection volume was again 5 µL, with mobile phase A consisting of 70% water + 5 mM ammonium formate (pH 4.0) and mobile phase B composed of 95% acetonitrile + 5 mM ammonium formate (pH 4.0). The flow rate was set to 0.25–0.4 mL/min, with the gradient program as follows: 0 minutes, 100% B (flow rate 0.25 mL/min); 1 minute, 100% B (flow rate 0.25 mL/min); 10.5 minutes, 60% B (flow rate 0.25 mL/min); 13 minutes, 15% B (flow rate 0.25 mL/min); 13.5 minutes, 15% B (flow rate 0.25 mL/min); 14 minutes, 100% B (flow rate 0.4 mL/min); 18.5 minutes, 100% B (flow rate 0.4 mL/min); 19 minutes, 100% B (flow rate 0.25 mL/min); 20 minutes, 100% B (flow rate 0.25 mL/min).

Targeted detection and quantification for plasma metabolites and lipids analysis was performed on an Agilent 6495 QqQ Mass Spectrometer using dynamic multiple reaction monitoring (dMRM) with an Agilent Jet Stream ion source. The polarity was switched between positive and negative modes. The gas temperature was set to 200 °C, with a drying gas flow rate of 14 L/min, nebulizer gas at 50 psi, and sheath gas at 375 °C and 12 L/min. The capillary voltage was set to 3,000 V for positive and –2,500 V for negative polarity, with a nozzle voltage of 0 V for both polarities. The iFunnel high/low pressure RF was set to 150/60 V for positive and 90/60 V for negative polarity. The scan type was set to dMRM with unit resolution for both Q1 and Q2, a delta EMV of 0 V (positive) and 200 V (negative), and a cell acceleration voltage of 5 V. The dMRM method was generated using Agilent MassHunter Acquisition software.

##### Development of Plasma Proteomics Fingerprint Database: CompleteBank-Discovered

For Plasma Proteins: We analyzed over 9,000 plasma proteomic runs using the timsTOF HT mass spectrometer (Billerica, Massachusetts, USA), systematically documenting the performance of each detected plasma protein. A stringent false discovery rate (FDR) threshold of 1% was applied in discovery-mode analysis to ensure high confidence in protein identification. By integrating data from these runs, we identified 17,328 unique plasma proteins. To facilitate validation on triple quadrupole (QqQ) mass spectrometers, we categorized these proteins into distinct classes based on their physicochemical properties and detection characteristics. A robust algorithm was developed to effectively translate protein fingerprints identified on the time-of-flight (TOF) platform to QqQ mass spectrometers. Validation assays were subsequently conducted using QqQ platforms from multiple vendors to confirm the reproducibility and reliability of these identified proteins.

Plasma Metabolites and Lipids: To establish a comprehensive plasma metabolite and lipid profile, we compiled a curated list of nearly 3,000 small molecules, including metabolites and lipids, based on an extensive literature review. Each molecule was subjected to at least six different analytical approaches to determine the optimal detection conditions in human plasma samples. To maximize the number of detectable small molecules while minimizing assay run time and improving throughput, we manually curated and optimized the resulting data. This refinement process led to the development of a high-efficiency detection protocol, ultimately documenting 2,927 molecules in our CompleteBank-Discovery database.

### Establishment of Validated and Optimized Detection Assay clusters: CompleteBank-Validated

For Plasma Proteins: To transition from discovery to validated clinical applications, we evaluated the clinical relevance of each identified plasma protein and selected an initial panel of 12,892 proteins from the discovery cohort. Extensive QqQ-based method optimization was performed for these proteins, involving iterative assays to fine-tune detection conditions, enhance sensitivity, and ensure reproducibility. Through this rigorous process, we refined the panel to a final set of **10,598 validated proteins**, optimized for reliable quantification using QqQ mass spectrometry.

#### Plasma Metabolites and Lipids

For targeted metabolomics and lipidomics, we established validated detection parameters for 762 metabolites and 1,395 lipids. To achieve optimal characterization, we employed three different chromatographic columns and implemented six distinct analytical methods, ensuring comprehensive coverage and precise quantification of these molecules.

### CompletePeaking: An Automatic Bioinformatic Pipeline for Clinical Proteomics and Metabolomics Data Processing

#### For Plasma Proteins

One of the essential parts of the *Complete360* pipeline is its methodology for peak picking and data analysis pipeline, and we term this entire bioinformatic pipeline as **CompletePeaking**. The CompletePeaking process combines manual curation with automated machine learning approaches to ensure the accurate identification and quantification of peptides in complex proteomic datasets. Initially, we employed the *Complete360* methods for manual validation of proteomics data, where human evaluators examined chromatograms and spectra for peptide transitions. The goal was to ensure that the peaks of interest coeluted with reference transitions and exhibited minimal background noise. Manual validation focused on assessing coelution of peaks and verifying that they exhibited high library dot product scores, which quantify the similarity between observed and reference intensity profiles.

Each peptide transition was scrutinized for quality indicators, including signal-to-noise ratio (SNR), intensity, and the presence of coelution, with a tolerance threshold of 0.3 minutes for retention time, as well as surrounding noise signals for each specific transitions. The manual curation process, although labor-intensive, was crucial in establishing the initial training dataset across diverse clinical samples, including several advanced stage cancer plasma samples, cardiovascular disease samples, neurodegenerative disease plasma samples and also inflammation and auto-immune disease plasma samples. These curated datasets formed the foundation for subsequent **CompletePeaking** model training.

#### Validation Process

During the manual validation phase, evaluators closely monitored the retention times of peptide transitions to ensure they coeluted. The measured retention times were then compared to predicted peak retention times, ensuring they fell within a defined tolerance. Dot product calculations were performed to assess the similarity between the observed and reference spectra, with high dot product scores indicating a strong match to the expected peptide identity. Peaks exhibiting both strong coelution and high dot product scores were labeled as "good" peaks, which were subsequently used as input for model training.

Despite the advantages of manual validation, it introduces certain biases. Evaluators often tend to prioritize peaks with higher intensity, which, while visually striking, may not always correspond to biologically relevant transitions. To mitigate these biases, we incorporated several strategies to improve the reliability of the training data. These included matching retention times across multiple samples, identifying reproducible noise landscapes around target peaks, and closely examining the patterns of transition similarity to the reference library. This comprehensive approach helped to ensure more accurate and consistent peak selection, enhancing the reliability of the training data.

A critical aspect of this study is the long-term accumulation of data, with years of manual peak validation forming the core of our training dataset. By manually curating and validating a large number of peaks across diverse clinical samples, we developed a robust dataset that captured the nuances of peak coelution, retention times, and library dot product scores. This curated dataset served as the backbone for training machine learning models, enabling the automation of peak selection while preserving the high accuracy and reliability achieved through manual methods.

#### Automated Peak Picking Using Machine Learning

To address the limitations of manual peak selection and improve scalability, we integrated an artificial intelligence-based machine learning model to automate the peak picking process. The model was trained using the XGBoost algorithm with a dataset that had been manually validated, incorporating various features derived from chromatogram analysis, such as coelution count, signal-to-noise ratio (SNR), shape correlation, and intensity. These features were essential for distinguishing high-quality peaks from those impacted by noise or background interference. A particularly important feature was coelution count, which represents the number of transitions that coelute at consistent retention times. A higher coelution count significantly boosts the confidence in identifying a valid peak. Additional features, including SNR, dot product, and shape correlation, were also considered, with their weights adjusted according to their contribution to the model’s overall performance. The model was trained using cross-validation to optimize hyperparameters and prevent overfitting, ensuring that it remained robust across different datasets. This process allowed the model to generalize well, improving its ability to reliably identify peaks across a variety of samples and conditions.

This combination of manual curation and automated machine learning not only enhances the accuracy and scalability of the peak picking process but also overcomes the limitations of traditional proteomics workflows. By integrating manual validation with machine learning models trained on years of curated data, we provide a more consistent, reliable, and high-throughput solution for proteomics studies. This ensures the identification of high-quality peaks across a variety of clinical samples, facilitating the analysis of large, complex datasets.

#### Data Preprocessing and Feature Generation

The preprocessing pipeline utilized CompleteBank-Discovered results to generate feature files, which provided a comprehensive list of candidate peaks based on observed chromatogram data. These candidate peaks were subsequently labeled according to manual validation results, with "good" peaks identified as those that met the quality criteria of coelution and high dot product scores. Features extracted from these peaks—including coelution count, dot product, signal-to-noise ratio (SNR), and shape correlation—were used as input for the XGBoost model. The model was trained to perform binary classification, distinguishing peaks as either valid or invalid based on their predicted retention times and associated feature characteristics. The model’s performance in predicting accurate peak retention times and selecting the most reliable candidate peaks was evaluated using a dataset comprising over 15,000 peptides, manually validated across a variety of representative clinical samples, including pooled advanced disease plasma samples. This robust dataset ensured that the model was trained on diverse, real-world data, enhancing its accuracy and generalizability.

#### Peak Selection, Postprocessing, and Data Normalization

After model training, automated peak selection was applied to new, unseen data. The postprocessing steps included selecting the highest-scoring peaks for each peptide sequence, ensuring that the final selection consisted of the most reliable peaks with the highest likelihood of accurate identification. Following peak selection, data normalization was performed on the peak area data collected from each target analyte. Given the variety of detection methods and clustering based on peak intensities, hydrophobicities, retention times, and other factors, normalization was conducted for each cluster using separate methods. As a result, multiple reference points were used for normalization, tailored to each cluster of target analysis. This approach led to the development of the Multi-point Normalized Protein eXpression (**MNPX**) value, which represents the normalized expression for each protein. After normalization, the abundance of each protein across various samples could be compared, enabling the evaluation of its correlation with disease states and facilitating the identification of potential protein biomarkers. For the current study, the normalization factor for each protein was chosen as the median intensity of the biomarker’s intensity in that detection cluster. Further normalization can be updated to use stably expressed plasma proteins or disease specific normalizers but will subject to further evaluation and development with at least several hundred samples for each disease type and will be updated in a further study.

### For Plasma Metabolites and Lipids

In CompletePeaking pipeline, we also developed a set of metabolomics peak picking algorithms based on a second derivative method. This approach was used to identify peaks in metabolomics datasets by analyzing the intensity profiles that are changing over time.

1. **Smoothing the Intensity Signal** The raw intensity data was smoothed using the Savitzky-Golay filter to reduce noise while preserving peak shape. The smoothing was applied with a window length of 11 and a polynomial order of 3.
2. **Computing the Second Derivative** The second derivative of the smoothed intensity was computed to capture the changes in concavity that mark the boundaries of a peak. These inflection points were used to define the start and end points of each peak.
3. **Zero-Crossings for Boundary Detection** Zero-crossings of the second derivative were identified, marking the transition from concave up to concave down. These zero-crossings defined the start and end time points of each peak. Adjustments were made to the boundaries to account for peak asymmetry, with an extension factor applied to both the start and end points.
4. **Background Correction and Tail Adjustment** A background intensity threshold was calculated as the median intensity outside the peak region. The peak boundaries were iteratively adjusted to ensure that they did not extend into noise regions, and the tail symmetry condition was met to balance the peak shape.
5. **Width Correction** To ensure consistent peak-width estimation across samples, outliers in peak width were removed using the interquartile range (IQR) method. After removing outliers, the peak widths were standardized based on the sample with the largest total area.

This second derivative method provided a robust framework for metabolomics peak detection, and it was incorporated as an essential part of CompletePeaking pipeline to identify and quantify metabolomic signals.

### Detectability in *Complete360* Methods

The *Complete360* platform employs a rigorous set of criteria to ensure high-confidence detection and quantification of target proteins. These criteria were established to maximize sensitivity, reproducibility, and specificity in deep proteomic profiling. The following parameters define detectability within the *Complete360* workflow:

1. **Signal-to-Noise Ratio (S/N):** Reliable detection of analytes requires a minimum signal-to-noise (S/N) ratio of 1.5, ensuring adequate signal intensity above background fluctuations.
2. **Retention Time (RT) Consistency:** Retention time (RT) for analyte detection must exceed 1.5 minutes, preventing interference from early eluting compounds and maintaining consistency across runs.
3. **Background and Interference Control:** To ensure specificity, analyte peaks must exhibit minimal interference from co-eluting species. Background signals within the retention window are required to be at least one order of magnitude lower than the analyte peak intensity. For example, an analyte peak with an intensity of 1,000 counts must have a background signal of ≤ 100 counts.
4. **Co-Elution of Fragment Ions:** The apexes of monitored transitions must closely align within a time window of ±0.05–0.2 minutes, ensuring the consistency of elution profiles and preventing misidentification.
5. **Fragment Ion Ratio Consistency:** Ion ratios between transitions of the same analyte must remain within ±20–30% of reference values, ensuring stability in detection across different analytical runs.
6. **Spectral Similarity Assessment:** To validate fragmentation patterns, spectral comparisons with high-resolution discovery data are performed. A dot-product or similarity score of ≥ 0.5–0.6 is required to confirm alignment with expected fragmentation profiles.

1. **Separation of high- and low-abundance molecules**: High- and low-abundance molecules should be detected separately to minimize ion suppression effects on low-abundance targets. This ensures that low-abundance molecules are detected with enhanced sensitivity and reproducibility, avoiding interference from high-abundance molecules, which typically overshadow other target analytes^23^.
2. **Resolution of co-eluted targets**: Co-eluted targets should be resolved at higher resolution when their abundance permits. While higher resolution may reduce the number of ions entering the mass analyzer of a mass spectrometer, potentially affecting sensitivity, careful manual tuning and optimization for each protein can address this.
3. **Retention time reproducibility**: Retention times for various analytes must be consistently reproducible across different assay environments. For instance, an analyte detected in a low pH fraction may exhibit a significantly different retention time compared to its detection in a high pH fraction. This variation underscores the importance of considering the surrounding microenvironment to ensure reproducible detection schedule.
4. **Management of co-eluting noise analytes**: High-abundance noise analytes must be carefully controlled to avoid co-elution with low-abundance target analytes. In our assay, some high-abundance analytes have been utilized to serve as highly reproducible landmarks, which can enhance data annotation accuracy and improve overall analytical precision which is further illustrated later.
5. **Minimizing Run Time**: To integrate all target analyses into a single assay with minimal runtime, an AI-driven pattern-clustering approach was employed. This method reduced overall run time while optimizing the sensitivity and reproducibility of each analyte when consolidated into the same assay. This AI-driven algorithm is an essential component of our in-house developed software package, which can also be used to establish customized diagnostic methods, such as those focused on specific diseases.

These criteria collectively ensure the robustness and accuracy of the *Complete360* methodology, facilitating the precise detection of ultra-low abundance proteins in complex biological matrices.

## Data Deposition

The data reported in this article have been deposited via ProteomeXchange in the PeptideAtlas SRM Experiment Library (PASSEL) (identifier PASS05916).

## Competing Interests Statement

All authors are current employees of Complete Omics Inc., the company that developed the Complete360 platform described in this study. As such, the authors have financial and professional interests related to the technologies and findings presented. This potential conflict of interest has been disclosed, and the study was conducted with a commitment to scientific rigor and integrity.

## Reference

1. Nicholson, B.D. et al. Multi-cancer early detection test in symptomatic patients referred for cancer investigation in England and Wales (SYMPLIFY): a large-scale, observational cohort study. Lancet Oncol 24, 733–743 (2023).

2. Klein, E.A. et al. Clinical validation of a targeted methylation-based multi-cancer early detection test using an independent validation set. Ann Oncol 32, 1167–1177 (2021).

3. Cohen, J.D. et al. Detection and localization of surgically resectable cancers with a multi-analyte blood test. Science 359, 926–930 (2018).

4. Chen, M. & Zhao, H. Next-generation sequencing in liquid biopsy: cancer screening and early detection. Hum Genomics 13, 34 (2019).

5. Davies, M.P.A. et al. Plasma protein biomarkers for early prediction of lung cancer. EBioMedicine 93, 104686 (2023).

6. Wang, Q. et al. Selected reaction monitoring approach for validating peptide biomarkers. Proc Natl Acad Sci U S A 114, 13519–13524 (2017).

7. FDA (2024).

8. Vitko, D. et al. timsTOF HT Improves Protein Identification and Quantitative Reproducibility for Deep Unbiased Plasma Protein Biomarker Discovery. J Proteome Res 23, 929–938 (2024).

9. Heil, L.R. et al. Evaluating the Performance of the Astral Mass Analyzer for Quantitative Proteomics Using Data-Independent Acquisition. J Proteome Res 22, 3290–3300 (2023).

10. Bonaventura, P. et al. Identification of shared tumor epitopes from endogenous retroviruses inducing high-avidity cytotoxic T cells for cancer immunotherapy. Sci Adv 8, eabj3671 (2022).

11. Hsiue, E.H. et al. Targeting a neoantigen derived from a common TP53 mutation. Science 371 (2021).

12. Terai, Y.L. et al. Valid-NEO: A Multi-Omics Platform for Neoantigen Detection and Quantification from Limited Clinical Samples. Cancers (Basel) 14 (2022).

13. Wang, Q. et al. Direct Detection and Quantification of Neoantigens. Cancer Immunol Res 7, 1748–1754 (2019).

14. Omenn, G.S., et al. The 2024 Report on the Human Proteome from the HUPO Human Proteome Project. J Proteome Res 23, 5296–5311 (2024).

15. Wishart, D.S. et al. MarkerDB: an online database of molecular biomarkers. Nucleic Acids Res 49, D1259–D1267 (2021).

16. Schooneveldt, Y.L. et al. The Impact of Simvastatin on Lipidomic Markers of Cardiovascular Risk in Human Liver Cells Is Secondary to the Modulation of Intracellular Cholesterol. Metabolites 11 (2021).

17. Su, B. et al. A DMS Shotgun Lipidomics Workflow Application to Facilitate High-Throughput, Comprehensive Lipidomics. J Am Soc Mass Spectrom 32, 2655–2663 (2021).

18. Cao, Z. et al. Evaluation of the Performance of Lipidyzer Platform and Its Application in the Lipidomics Analysis in Mouse Heart and Liver. J Proteome Res 19, 2742–2749 (2020).

19. Medina, J. et al. Omic-Scale High-Throughput Quantitative LC-MS/MS Approach for Circulatory Lipid Phenotyping in Clinical Research. Anal Chem 95, 3168–3179 (2023).

20. Tianqi Chen, C.G. XGBoost: A Scalable Tree Boosting System. arXiv:1603.02754 (2016).

21. Ignjatovic, V. et al. Mass Spectrometry-Based Plasma Proteomics: Considerations from Sample Collection to Achieving Translational Data. J Proteome Res 18, 4085–4097 (2019).

22. Millioni, R. et al. High abundance proteins depletion vs low abundance proteins enrichment: comparison of methods to reduce the plasma proteome complexity. PLoS One 6, e19603 (2011).

23. Tu, C. et al. Depletion of abundant plasma proteins and limitations of plasma proteomics. J Proteome Res 9, 4982–4991 (2010).

24. Douglass, J., et al. Bispecific antibodies targeting mutant RAS neoantigens. Sci Immunol 6 (2021).

25. Deutsch, E.W. et al. Advances and Utility of the Human Plasma Proteome. J Proteome Res 20, 5241–5263 (2021).

26. Pettersson, F. et al. Ribavirin treatment effects on breast cancers overexpressing eIF4E, a biomarker with prognostic specificity for luminal B-type breast cancer. Clin Cancer Res 17, 2874–2884 (2011).

27. Gao, Y. et al. Loss of ERalpha induces amoeboid-like migration of breast cancer cells by downregulating vinculin. Nat Commun 8, 14483 (2017).

28. Ding, Q. et al. Ficolin-2 triggers antitumor effect by activating macrophages and CD8(+) T cells. Clin Immunol 183, 145–157 (2017).

29. Cao, B. et al. Cancer-mutated ribosome protein L22 (RPL22/eL22) suppresses cancer cell survival by blocking p53-MDM2 circuit. Oncotarget 8, 90651–90661 (2017).

30. Niu, L. et al. Plasma proteome variation and its genetic determinants in children and adolescents. Nat Genet (2025).

31. Obradovic, M., et al. Leptin and Obesity: Role and Clinical Implication. Front Endocrinol (Lausanne) 12, 585887 (2021).

32. Candia, J. et al. Assessment of Variability in the SOMAscan Assay. Sci Rep 7, 14248 (2017).

33. Candia, J., Daya, G.N., Tanaka, T., Ferrucci, L. & Walker, K.A. Assessment of variability in the plasma 7k SomaScan proteomics assay. Sci Rep 12, 17147 (2022).

34. Smits, H.M. et al. The BAMBOO method for correcting batch effects in high throughput proximity extension assays for proteomic studies. Sci Rep 15, 1498 (2025).

35. Dennis, M.S. et al. Albumin binding as a general strategy for improving the pharmacokinetics of proteins. J Biol Chem 277, 35035–35043 (2002).

36. Hui, H., Farilla, L., Merkel, P. & Perfetti, R. The short half-life of glucagon-like peptide-1 in plasma does not reflect its long-lasting beneficial effects. Eur J Endocrinol 146, 863–869 (2002).

37. Razavi, M. et al. Measuring the Turnover Rate of Clinically Important Plasma Proteins using an Automated SISCAPA Workflow. Clin Chem 65, 492–494 (2019).

38. Wang, Y. et al. Reversed-phase chromatography with multiple fraction concatenation strategy for proteome profiling of human MCF10A cells. Proteomics 11, 2019–2026 (2011).

39. Demichev, V., Messner, C.B., Vernardis, S.I., Lilley, K.S. & Ralser, M. DIA-NN: neural networks and interference correction enable deep proteome coverage in high throughput. Nat Methods 17, 41–44 (2020).

40. Kong, A.T., Leprevost, F.V., Avtonomov, D.M., Mellacheruvu, D. & Nesvizhskii, A.I. MSFragger: ultrafast and comprehensive peptide identification in mass spectrometry-based proteomics. Nat Methods 14, 513–520 (2017).

